# Molecular basis of human asparagine synthetase inhibitor specificity

**DOI:** 10.1101/428508

**Authors:** Wen Zhu, Ashish Radadiya, Claudine Bisson, Brian E. Nordin, Patrick Baumann, Tsuyoshi Imasaki, Sabine A. Wenzel, Svetlana E. Sedelnikova, Alexandria H. Berry, Tyzoon K. Nomanbhoy, John W. Kozarich, Yi Jin, Yuichiro Takagi, David W. Rice, Nigel G. J. Richards

## Abstract

Expression of the enzyme human asparagine synthetase (ASNS) promotes metastatic progression in breast cancer, which affects L-asparagine levels and tumor cell invasiveness. Human ASNS has therefore emerged as a *bona fide* drug target for cancer therapy. We have reported a slow-onset, tight binding ASNS inhibitor with nanomolar affinity, but our compound exhibits poor cell permeability. On the other hand, we show here that this inhibitor exhibits remarkable selectivity for the human ASNS in HCT-116 cell lysates. By determining the first high-resolution (1.85 Å) X-ray crystal structure for human ASNS, we have built a computational model of the enzyme complexed to our inhibitor, which provides the first insights into the intermolecular interactions mediating specificity. These findings should facilitate the development of a second generation of ASNS inhibitors, leading to the discovery of drugs to prevent metastasis.

---

Asparagine synthetase (ASNS) catalyzes the ATP-dependent biosynthesis of L-asparagine in cells from L-aspartic acid using L-glutamine as a nitrogen source^1^. L-asparagine is important for the growth and maintenance of acute lymphoblastic leukemias^2^, and breast^3^, lung^4^ and castration-resistant prostate^5^ cancers. Moreover, altering exogenous L-asparagine levels affects tumor cell invasiveness, and enforced expression of human ASNS promotes metastatic progression in breast cancer^6^, presumably by increasing the bioavailability of L-asparagine. Consistent with these findings, functional genomic screening of a murine sarcoma model, generated by oncogenic forms of *Kras*^7,8^, has demonstrated that silencing the gene encoding ASNS inhibits cell proliferation^9^. All of these observations strongly suggest that human ASNS is a *bona fide* drug target and that potent, small molecule ASNS inhibitors could have significant clinical utility in the prevention of metastasis^1^, and perhaps more broadly in cancer chemotherapy^10^.

The need to develop ASNS inhibitors was recognized over 30 years ago. With the recent emergence of human ASNS as a drug target for cancer therapy, this need has grown significantly. Identifying compounds with nanomolar affinity for the enzyme, however, has proved to be remarkably difficult. The only reported screening studies failed to identify small molecules with sub-micromolar binding and/or high selectivity for human ASNS^11,12^, probably because of a lack of mechanistic and structural information about the enzyme. We therefore elucidated the kinetic and catalytic mechanisms of the glutamine-dependent asparagine synthetase (AS-B)^13–15^ encoded by the *asnB* gene in *Escherichia coli*^16^. These studies revealed that both the β-aspartyl-AMP intermediate and the transition state for its subsequent reaction with ammonia were tightly bound by the enzyme during catalysis (**Fig. 1a**)^14^. Although unreactive analogs of the β-aspartyl-AMP intermediate are sub-micromolar ASNS inhibitors^17^, the recognition that methylsulfoximines mimic the key transition state for the attack of ammonia on activated esters^18–20^ led to the synthesis of the functionalized methylsulfoximines **1** and **2** (**Fig. 1b**), which are slow-onset inhibitors that exhibit nanomolar affinity for the enzyme^21,22^.

**Fig. 1:**
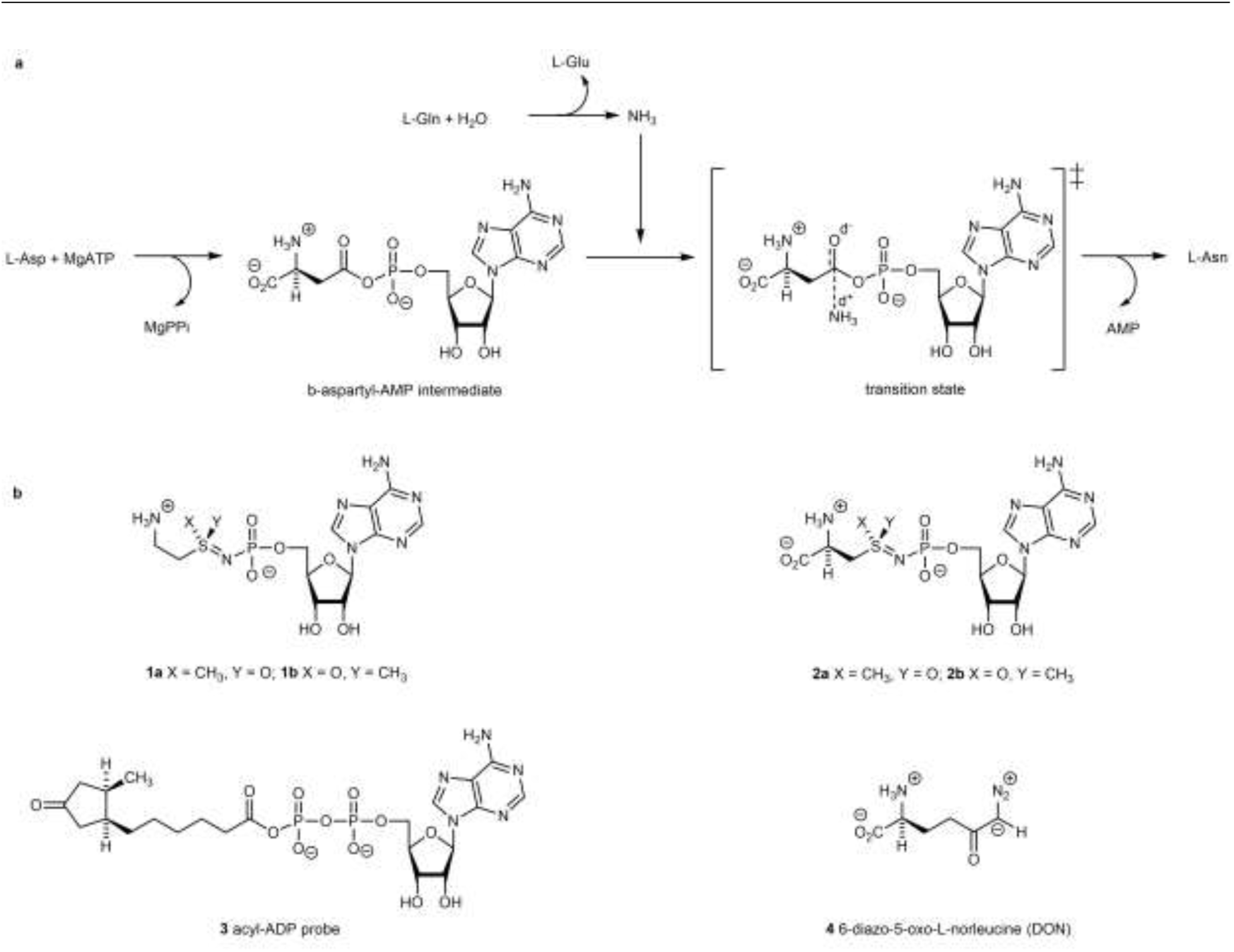
Catalytic mechanism of human ASNS and structures of compounds 1–4. (**a**) Overview of the chemical transformations catalyzed by ASNS showing the β-aspartyl-AMP intermediate and the transition state for its subsequent reaction with an ammonia molecule, which is released from L-glutamine in a separate glutaminase site. (**b**) Chemical structures for functionalized methylsulfoximines **1** and **2**, activity-based probe **3** and 6-diazo-5-oxo-L-norleucine **4**.

Unfortunately, both of these compounds can only be prepared as a 1:1 mixture of diastereoisomers (**Fig. 1b**) that cannot be separated on column chromatography, severely limiting their usefulness in studies employing animal models of cancer and metastasis. In addition, the presence of the charged functional groups in the methylsulfoximines **1** and **2** likely contribute to poor cell permeability. Nonetheless, and very importantly, ASNS inhibitor **1** (**Fig. 1b**) does negatively impact the growth of sarcoma cells in a manner similar to that seen when ASNS expression is reduced using siRNA knockdown methods^9^. Moreover, this compound is cytotoxic against asparaginase-resistant MOLT-4 leukemia cells when used at micromolar concentrations^21^. Thus, ASNS inhibitor **1** does possess anti-cancer properties. We now report that **1** exhibits a remarkable degree of specificity against human ASNS in cell lysates, suggesting that this compound can serve as a starting point for drug discovery. Developing an in-depth understanding of the molecular basis for this unique specificity is absolutely required, however, if efforts to generate a second generation of small molecule ASNS inhibitors, which have improved bioavailability and lower chemical complexity, are to be successful.

## Results

### Evaluating the binding specificity of ASNS inhibitor 1 in human cell lysates

We undertook functionalized proteomics experiments employing the chemically reactive probe **3** (**Fig. 1**)^23^, which has been used to determine inhibitor selectivity against native kinases or other ATPases by profiling compounds in lysates derived from cells or tissues (KiNativ)^24^, to evaluate the affinity of this potent ASNS inhibitor **1** for alternate targets with a focus on non-kinase ATPases. We incubated probe molecule **3** with HCT-116 cell lysates in the presence and absence of the ASNS inhibitor. HCT-116 cells can metastasize in xenograft models and have been used in studies of colon cancer proliferation^25^. MS/MS fragmentation and sequence analysis of the tryptic peptides obtained from the two reaction mixtures showed that ASNS inhibitor **1** suppressed the ability of the probe to acylate the side chain of Lys-466 (located within the ATP-binding site of human ASNS) to an extent of 62% when present in cell lysates at 10 μM total concentration (**Table 1 and Supplementary Table 1**). As importantly, only moderate to low suppression of lysine acylation by the reactive probe **3** was seen for potential “off-target” enzymes when the ASNS inhibitor was present in the HCT-116 lysates. For example, ASNS inhibitor **1** did not suppress lysine acylation in the ATP-binding sites of GMP synthetase^26^, argininosuccinate synthetase^27^, and glutamyl-tRNA synthetase^28^ at 10 μM concentration even though these three enzymes convert ATP to AMP and PP_i_ during catalytic turnover. Moreover, the ASNS inhibitor **1** exhibited only weak binding to phosphopantetheine adenylyltransferase^29^ and nicotinate-nucleotide adenylyltransferase^30^ in the cell lysates at 10 μM concentration (**Table 1**). We also examined the ability of the ASNS inhibitor **1** to interact with the adenylylation sites of amino-acyl tRNA synthetases because the structurally similar compound **2** (**Fig. 1b**) was shown to bind tightly to *Escherichia coli* ammonia-dependent asparagine synthetase (AS-A)^31^, which is evolutionarily related to this class of enzymes^32^. Our experiments confirm that ASNS inhibitor **1** does not interact with lysyl, seryl or asparaginyl tRNA synthetases when present in the lysate at 10 μM concentration (**Table 1**). This observation is remarkable given that the related compound **2** (**Fig. 1b**) inhibits the *Escherichia coli* ortholog of asparaginyl-tRNA synthetase^31^. Increasing the concentration of the inhibitor **1** to 100 μM in the cell lysate suppressed Lys-466 acylation by the reactive probe **3** to an extent of 73%, a value that is unexpectedly low given that *in vitro* kinetic assays show that the ASNS inhibitor **1** inhibits recombinant, human ASNS with a nanomolar 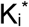 value^21^. Several explanations might account for this finding. For example, the reactive probe **3** might successfully compete for the free enzyme in the initial phase of incubation because the ASNS inhibitor **1** exhibits slow-onset, tight-binding kinetics. The ability of **1** to bind to the synthetase site may also be negatively impacted by the ATP concentration in the cell lysates, which is in the range of 1–10 mM. Even when present at a concentration of 100 μM, however, ASNS inhibitor **1** still exhibits remarkable selectivity (**Table 1 and Supplementary Table 1**) and the apparent absence of any interaction with kinases suggests that the development of highly specific ASNS inhibitors with more “drug-like” chemical structures^33^ can be achieved.

**Table 1.**
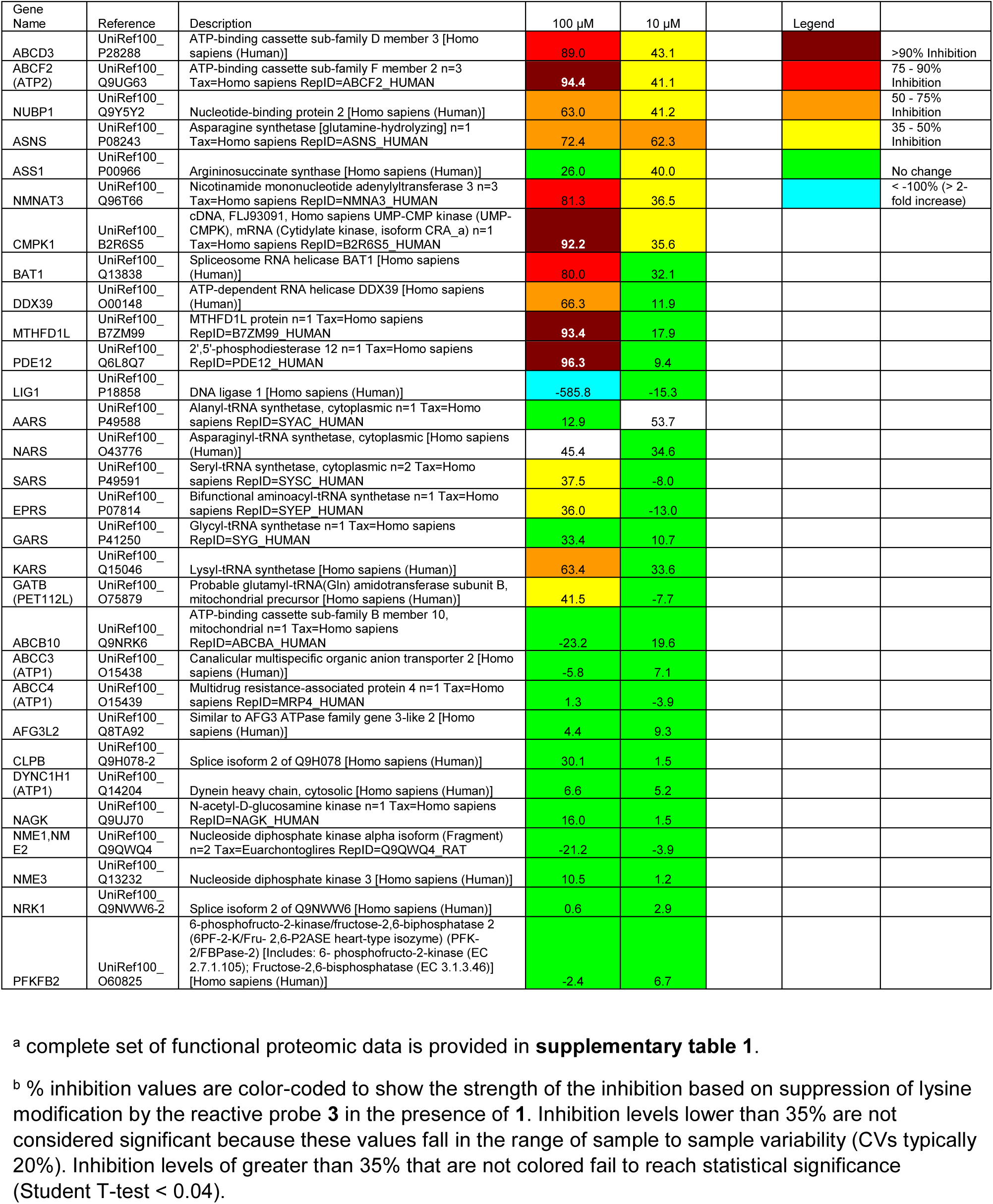
Selected results from the functional proteomics assay showing the % inhibition of cellular ATPases by the ASNS inhibitor 1 at 100 μM and 10 μM concentration.^a,b^

### Molecular structure of human ASNS

In order to determine the molecular basis for the observed specificity of ASNS inhibitor **1**, and to provide a firm basis for identifying simplified molecular scaffolds for potent and selective ASNS inhibitors, we obtained the first high resolution X-ray crystal structure of human ASNS. To date, only the structure of the glutamine-dependent ASNS (AS-B) encoded by the *asnB* gene in *Escherichia coli* has been reported^34^, and efforts to obtain crystals of this ASNS homolog bound to small molecules other than AMP have proven unsuccessful. Moreover, differences in the ability of the functionalized methylsulfoximine **2** (as a 1:1 mixture of the diastereoisomers **2a** and **2b**)(**Fig. 1b**) to inhibit the human and bacterial forms of the enzyme have been reported^22,35^. Multi-milligram amounts of highly active, recombinant, C-terminally His10-tagged, human ASNS were obtained by expression in Sf9 cells^36^ by optimizing its expression using TEQC method as described (**see Supplementary Information**).^37^ The enzyme was initially purified by metal-affinity chromatography followed by removal of the C-terminal His10-tag by digestion with the S219P variant of TEV protease^38^. The resulting sample of untagged ASNS was then reacted with DON (6-diazo-5-oxo-L-norleucine)^39^ **4** (**Fig. 1a**) to modify the reactive thiolate of Cys-1 in the glutaminase active site of ASNS^40^. Under our reaction conditions, the DON-modified form of human ASNS was obtained as a homogeneous protein in which other cysteine residues in the protein were not modified on the basis of mass spectrometry analysis of the “as-purified” and DON-modified protein (**Supplementary Fig. 1**). These mass spectrometric measurements also showed that the N-terminal methionine residue of the recombinant enzyme had been correctly processed to give the “mature” form of the enzyme. Conditions were then identified that gave single crystals of DON-modified human ASNS, one of which diffracted to 1.85 Å resolution (**Supplementary Table 2**). The structure of the enzyme was solved by molecular replacement^41^ using AS-B^34^ as a search model. Two molecules of DON-modified human ASNS were present in the asymmetric unit as a head-to-head dimer in which the two monomers were linked by a disulfide bond that likely formed during crystallization (**Fig. 2a and Supplementary Fig. 3**).

**Fig. 2:**
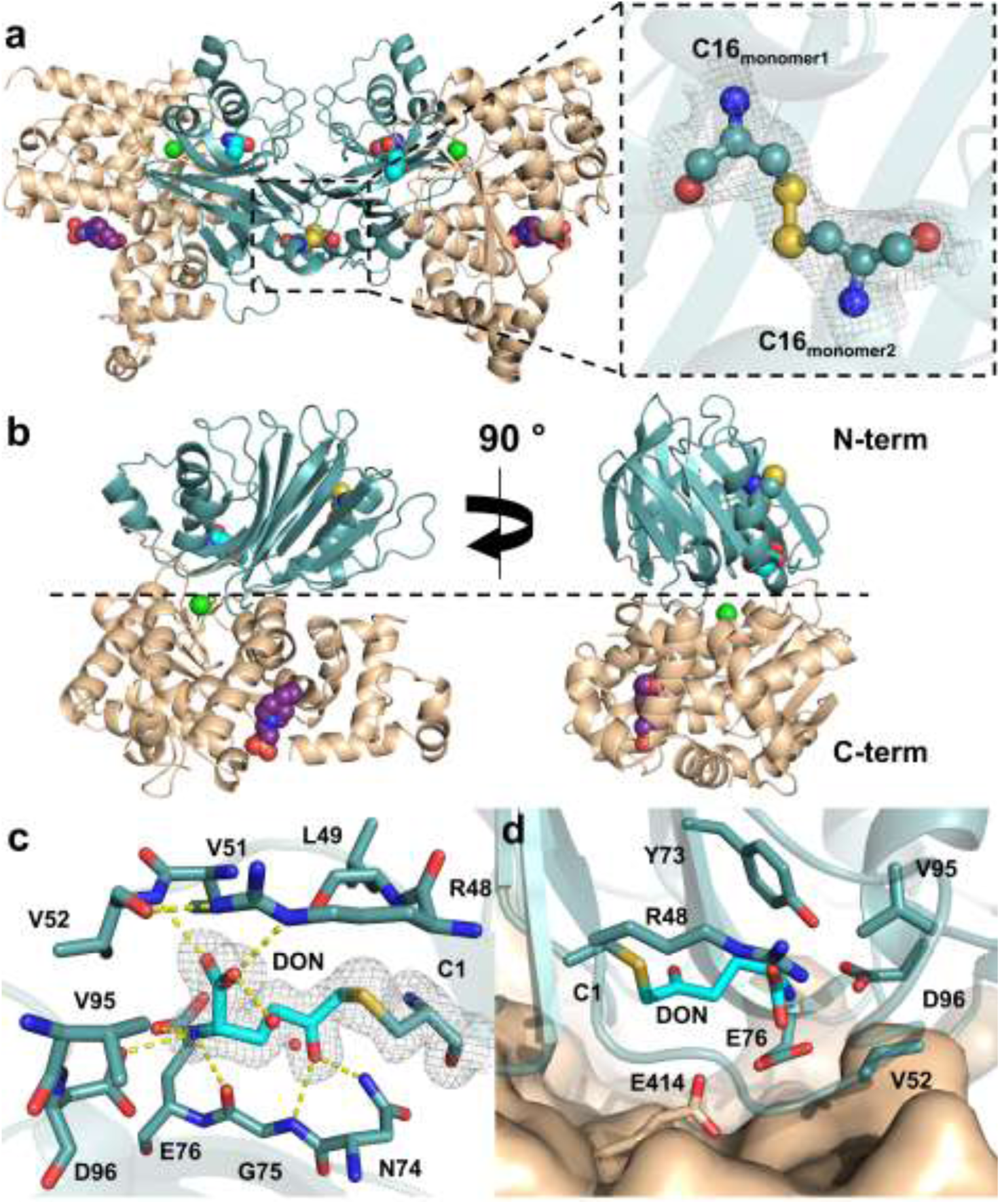
X-ray crystal structure of human ASNS and the glutaminase active site. (**a**) (Left) Cartoon representation of the asymmetric unit of the human ASNS crystal structure. The N-and C-terminal domains are colored blue and tan, respectively, and the atoms in the disulfide bond joining the monomers are rendered as spheres. Carbon atoms in the DON moiety and the HEPES molecule present in the synthetase active site are shown as cyan and purple spheres, respectively, and the bound chloride ion is drawn as a green sphere. (Right) Close-up of the disulfide bond connecting the N-terminal domains of the ASNS monomers and the electron density surrounding this region (grey mesh, contoured at 0.5σ). Color scheme: C, cyan; N, blue; O, red; S, yellow). (**b**) Cartoon representation of the human ASNS monomer. Domain and atom coloring scheme is identical to that used in (**a**). (**c**) Close-up of DON-modified Cys-1 showing the hydrogen bonding interactions between the DON moiety (cyan) and residues in the glutaminase active site. Electron density for DON, contoured at 0.5σ, is rendered as a mesh. (**d**) Location of the glutaminase site at the domain/domain interface of human ASNS showing the salt bridge involving the Arg-48 and Asp-96 side chains. Residues are identified using standard one-letter codes, and are numbered from the N-terminal residue (Cys-1).

As is seen for the bacterial homolog, human ASNS is composed of two domains (**Fig. 2b**). Residues in C-terminal (residues 203-560) synthetase domain (41.6% identity) are more conserved than those in the N-terminal (residues 1–202) glutaminase domain (33.9% identity) based on sequence comparisons of human ASNS and its homologs in a number of model organisms (**Supplementary Fig. 4**).

The N-terminal domain of human ASNS possesses the typical sandwich-like α/β/β/α topology seen in other N-terminal nucleophile (Ntn) amidotransferases^40,42^, such as GMP synthetase,^43^ and glutamine PRPP amidotransferase^44^. As seen in the structure of the bacterial homolog AS-B, a cis-proline (Pro-60) linkage is present in the human enzyme. Electron density for the DON-modified Cys-1 side chain is clearly evident in each monomer (**Fig. 2c**). A hydrogen bond network, composed of the conserved residues Arg-48, Val-52, Asn-74, Gly-75, Glu-76, and Asp-96, which mediates substrate recognition and thioester stabilization in the hydrolysis reaction that produces ammonia, is also clearly defined^45^. This substrate-binding pocket is located at the interface of the two domains, and is within 5 Å of an absolutely conserved glutamate residue (Glu-414) in the C-terminal domain (**Fig. 2d**). After refinement of the protein and ligands, a single 12 σ peak remained in a pocket at the interface of the N-and C-terminal domains on both chains. The site is surrounded by Tyr-78, Arg-416, Arg-245 and Val-417, and the peak was assigned as a chloride anion (**Fig. 2b**). The functional importance of this finding remains to be established for human ASNS, but plant asparagine synthetases are known to be activated by chloride.^46^ Importantly for structure-based inhibitor discovery, the synthetase site in the C-terminal domain, which is composed of sixteen α-helices and five β-strands, is well resolved (**Fig. 2b**). Unexpectedly, we observed a bound HEPES molecule from the crystallization buffer in the active site of this domain, which hydrogen bonds to the Asp-334 side chain and water molecules in a network that also involves conserved residues Asp-400 and Arg-403 (**Fig. 3a and 3b**). Residues in the synthetase active sites of human ASNS and AS-B are highly conserved (**Supplementary Fig. 4**) except that Val-272 and Met-333 in the bacterial form of the enzyme are replaced by Ile-287 and Ile-347, and superimposing the human and bacterial structures confirms that the active sites are almost identical (**Fig. 3c**). The Arg-403 side chain, however, adopts different conformations, presumably because AMP is not present in the human ASNS structure.

**Fig. 3:**
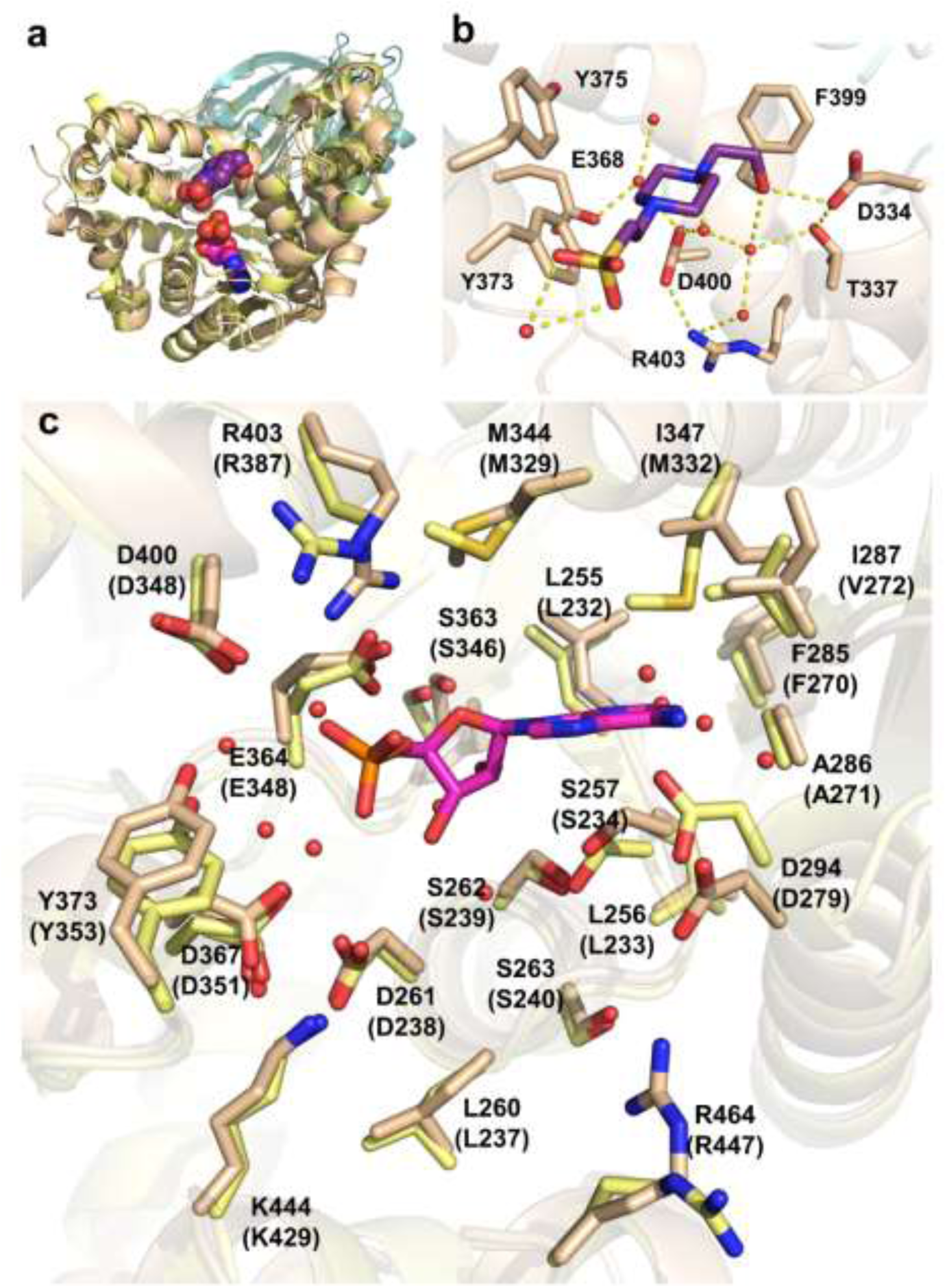
Structural features of the synthetase active site and putative inhibitor binding pocket. (**a**) Cartoon rendering of DON-modified human ASNS and the C1A AS-B variant (1CT9)^34^ containing a molecule of HEPES (C: purple spheres) and AMP (C: magenta spheres) in the two synthetase active sites, respectively. The N-and C-terminal domains of human ASNS are colored blue and tan, respectively, while the cognate domains in AS-B are colored green and pale yellow, respectively. (**b**) Close-up view of the network of intermolecular interactions between bound HEPES and residues/water molecules in the synthetase active site of human ASNS. Hydrogen bonds are shown as yellow dashed lines and waters by red spheres. (**c**) Residue conservation in the synthetase active sites of human ASNS and AS-B with side chain carbons of the two homologs being colored tan and pale yellow, respectively. Carbon atoms in the AMP molecule observed in the AS-B synthetase site are rendered in magenta. Waters are rendered as red spheres. Residues in both structures are identified using one-letter codes, and are numbered from the N-terminal residue (Cys-1). AS-B residue numbers are given in parentheses.

### Asparagine synthetase deficiency

Having the crystal structure of human ASNS in hand offered an opportunity to map the locations of mutations in 15 residues (Ala-5, Arg-48, Leu-144, Leu246, Gly-288, Thr-336, Arg-339, Phe-361, Ala-379, Tyr-397, Arg406, Ser-479, Val-488, Trp-540 and Arg-549) that have been identified in patients with asparagine synthetase deficiency (ASD) (**Fig. 4**)^47^. ASD is a rare neurological disorder that severely affects brain development; indeed, children with the disease often exhibit microcephaly, epileptic-like seizures and intellectual disability^48^. Three of these mutational locations are in the N-terminal domain although, of these, only the Arg-48 side chain is positioned such that it could interact directly with the L-glutamine substrate in the glutaminase active site. Substituting other amino acids in this position might therefore impact the chemical steps leading to ammonia formation. The remaining sites, which are mainly located in the C-terminal, synthetase domain can be clustered into four groups (**Supplementary Table 3**). Given that none of these residues seem to be positioned adjacent to L-aspartate or ATP in the synthetase active site, we hypothesize that mutations in these locations may perturb the structural stability and/or dynamic properties of the enzyme. Of course, the apparent involvement of ASNS in neurological development implies that clinically useful ASNS inhibitors must not cross the blood-brain barrier or be restricted to use in adults.

**Fig. 4:**
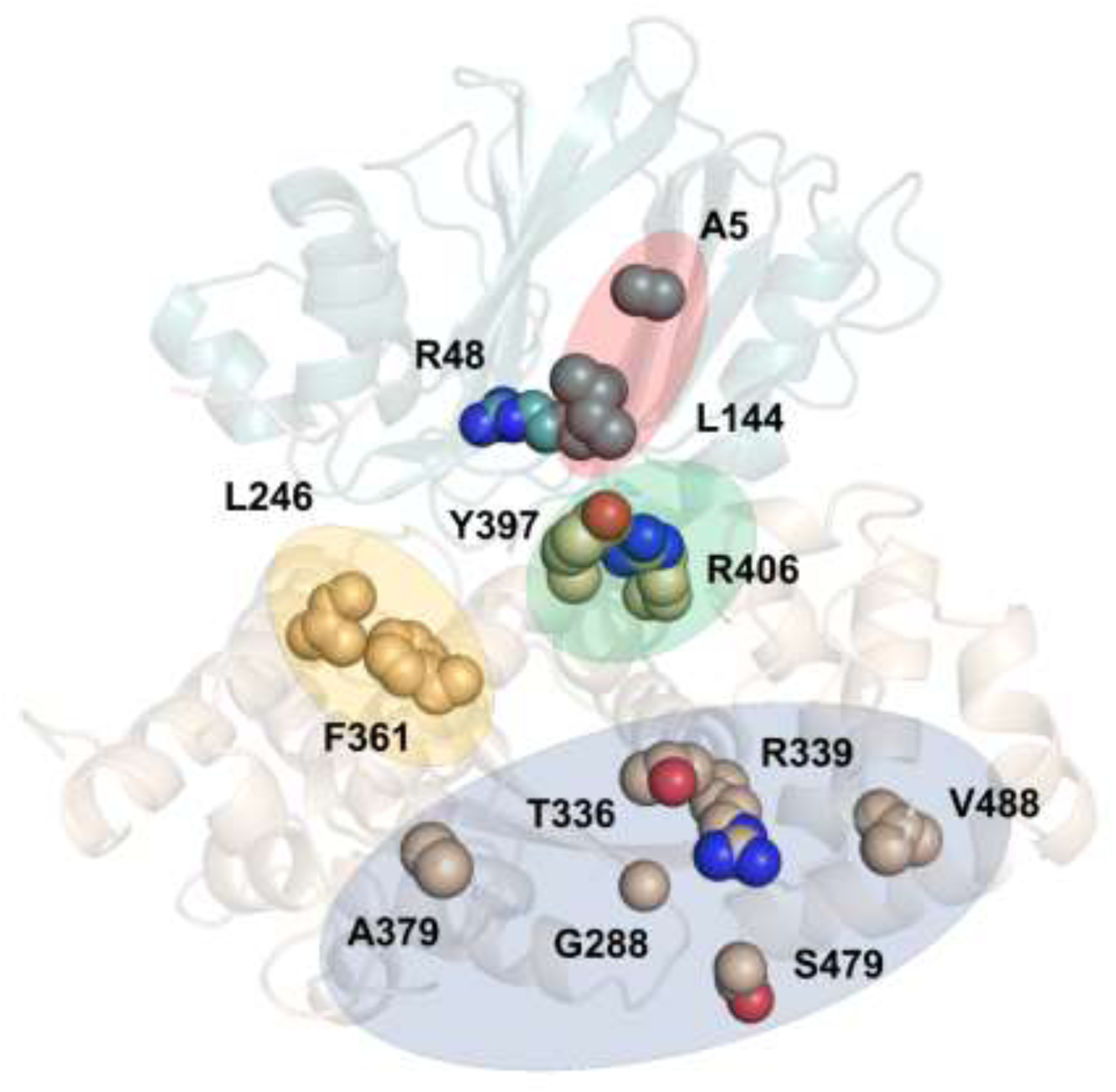
Locations of mutated residues in human ASNS that are associated with asparagine synthetase deficiency. (**a**) Cartoon representation of the X-ray crystal structure of human ASNS showing the locations of the 15 mutational sites (Ala-5, Arg-48, Leu-144, Leu246, Gly-288, Thr-336, Arg-339, Phe-361, Ala-379, Tyr-397, Arg-406, Ser-479, Val-488, Trp-540 and Arg-549) that have been identified in patients with asparagine synthetase deficiency. The side chains of mutated residues are rendered as spheres. Colored regions indicate sites that can be classified as Group 1 (red), Group 2 (yellow), Group 3 (green) and Group 4 (blue) (**Supplementary Table 3**).

### Structurally related human proteins

We sought to exploit our X-ray crystal structure in determining the molecular basis of the high binding selectivity exhibited by ASNS inhibitor **1** in the functional proteomics studies (**Table 1**). 128 proteins possessing domains with significant structural similarity to the ASNS synthetase domain (222–533) (**Supplementary Table 4**), of which 80 are not present in human cells were identified using the Dali server (http://ekhidna2.biocenter.helsinki.fi/dali/)^49^. Almost all of remaining 48 “structural neighbors” (uroporphyrinogen-III synthase^50^ being an interesting exception) employ ATP or structurally related molecules (SAM and NAD^+^) as substrates; X-ray crystal structures of the human homologs have been reported, however, for only 10 of these enzymes (**Supplementary Table 4**). Based on mechanistic considerations, ASNS inhibitor **1** might be expected to bind to glutamine-dependent NAD^+^ synthetase^26^, GMP synthetase^27^, argininosuccinate synthetase^51^ and FMN adenylyltransferase^52^, and, indeed, these enzymes have the highest Z-scores for proteins having AMP-forming domains homologous to the C-terminal domain of human ASNS. Moreover, all five of these enzymes possess a conserved sequence motif SGGxD (PP motif)^53^, which binds inorganic pyrophosphate produced concomitantly with the adenylylated intermediate during catalytic turnover. Unfortunately, tryptic peptides from the ATP-binding sites of glutamine-dependent NAD^+^ synthetase and FMN adenylyltransferase were not observed in the functional proteomics assay, suggesting that neither of these enzymes were present in HCT-116 cells under our growth conditions.

### Modeling the interaction of ASNS inhibitor 1 with human ASNS

Nevertheless, in order to understand why the transition state analog **1** does not bind to GMP synthetase or argininosuccinate synthetase at 10 μM concentration, we decided to obtain a structure for this inhibitor **1** bound to the human enzyme. Kinetic measurements show that **1** is competitive with respect to ATP (the first substrate to bind in the pathway leading to asparagine formation^14^), and can bind to the DON-modified form of human ASNS (data not shown). Both of these observations are consistent with the idea that the functionalized methylsulfoximine **1** binds within the synthetase active site of the enzyme (**Fig. 3**). Extensive crystallization trials, however, failed to yield crystals of ASNS inhibitor **1** bound to either DON-modified or wild type human ASNS. We therefore built computational models of the enzyme bound to each of the diastereoisomers **1a** and **1b** (**Fig. 5a**) in which Cys-1 was present as the unmodified amino acid.

**Fig. 5:**
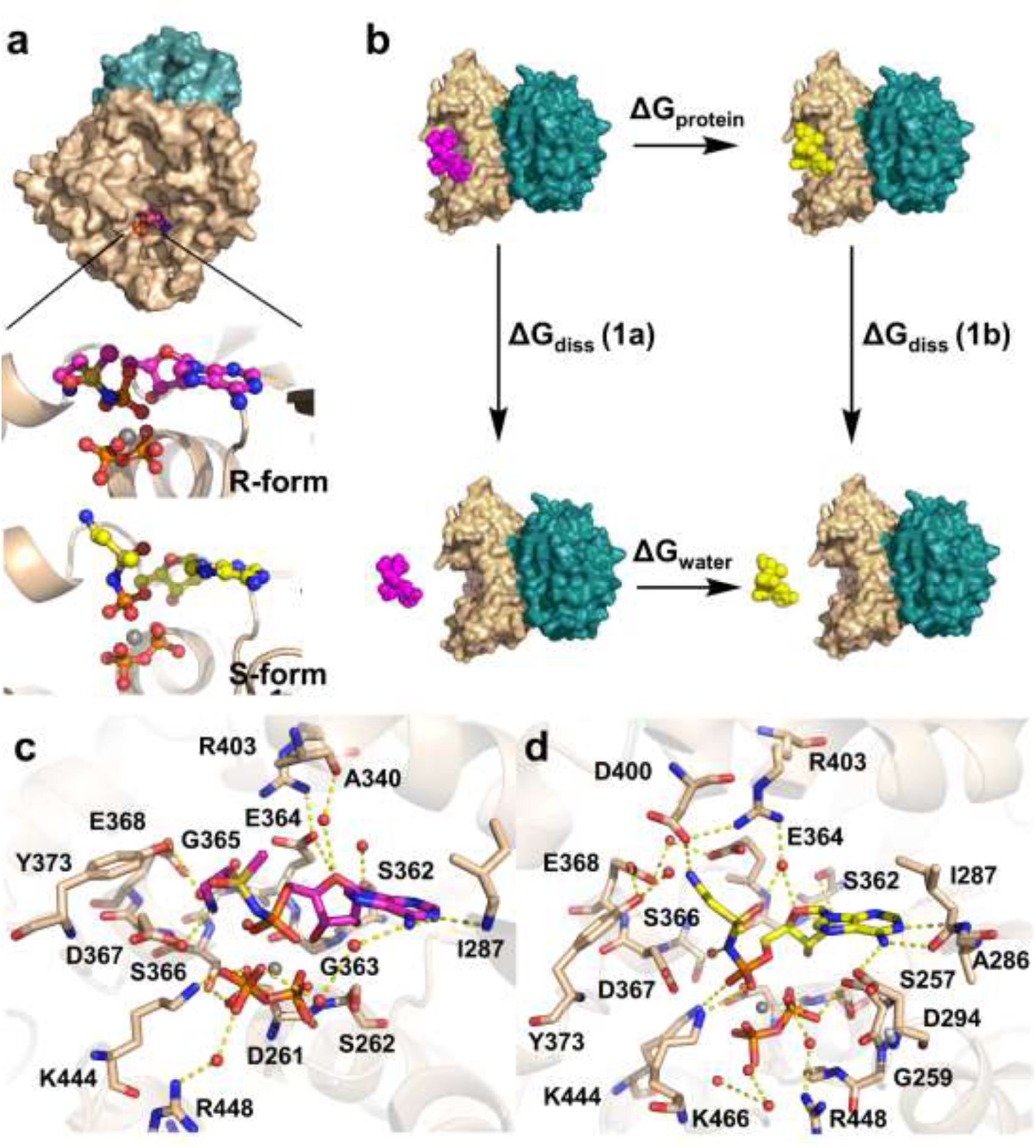
Computational models of the functionalized methylsulfoximines 1a and 1b bound within the synthetase active site of the human ASNS/MgPPi complex. (**a**) Surface representation of human ASNS showing the location of the putative inhibitor binding pocket within the synthetase active site. The N-and C-terminal domains of the enzyme are colored in blue and tan, respectively. Close-up views show the positions of the functionalized methylsulfoximines **1a** (green) and **1b** (yellow) relative to the bound inorganic pyrophosphate (orange) and Mg^2+^ (grey) ions in each model complex. (**b**) Thermodynamic cycle used to estimate the difference in binding free energy (ΔG_bind_ = ΔG_bind_(**1a**) −ΔG_bind_(**1b**)) of the diastereoisomers **1a** and **1b** computed from ΔG_protein_ – ΔG_water_ values obtained by free energy perturbation calculations. Both ΔG_bind_(**1a**) and ΔG_bind_(**1b**) have positive values since they describe dissociation of each ASNS/ligand complex. (**c**) Close-up of **1a** showing the non-covalent interactions with synthetase active site residues and water molecules in the computational model. (**d**) Close-up of **1b** showing the non-covalent interactions with synthetase active site residues and water molecules in the computational model.

We also included inorganic pyrophosphate (MgPP_i_) in the synthetase site because MgPP_i_ is the last product released during turnover^16^ and is therefore present in the enzyme when ammonia reacts with the β–aspartyl-AMP intermediate. MgPP_i_ was positioned above a conserved pyrophosphate-binding motif in a similar location to that seen in GMP synthetase^43^. All missing loops in the human ASNS structure, together with residues 534–546 located in the C-terminal tail, were built the procedures described in **Supplementary Information** with only the last fourteen residues (547–560) of the enzyme being omitted. *In silico* docking^54^ was used to position each of the diastereoisomers **1a** and **1b** into the MgPPi/ASNS complex, and the resulting models were placed in a box of water molecules. Molecular dynamics (MD) simulations (100 ns) were performed for both of the solvated model complexes (**see Supplementary Information**), with the free enzyme and the β–aspartyl-AMP/MgPPi/ASNS complex being used as “control” structures. We observe an extensive series of non-covalent interactions between β– aspartyl-AMP and the enzyme (**Supplementary Fig. 7a**). For example, the phosphate moiety of this reactive intermediate forms an electrostatic interaction with the conserved Lys-466 side chain (**Supplementary Fig. 4**), which has been shown to be essential for activity in AS-B^55^. In addition, the 2’-OH group on the ribose ring hydrogen bonds to the side chain of Ser-362, providing a molecular explanation for the fact that dATP is not a substrate. At the other end of the intermediate, the α-amino group forms a salt bridge with the conserved Asp-367 side chain and the α-carboxylate interacts with Glu-364 via a water molecule. To our knowledge, neither of these residues has been altered by site-directed mutagenesis even though both are conserved within known asparagine synthetases (**Supplementary Fig. 4**). A similar set of interactions was observed during the MD-derived trajectories of the model complexes containing ASNS inhibitor **1** (methylsulfoximines **1a** and **1b)**. Importantly for future inhibitor discovery efforts, the positively charged amino group of each functionalized methylsulfoximine is preferentially bound in a pocket defined by the side chains of Glu-364, Asp-367 and Asp-400 (**Fig. 5c**). Protein/ligand hydrogen bonds are also formed between both **1a** and **1b** and Ser-362, Gly-363 and Ile-287, and explicit waters mediate interactions between the functionalized methylsulfoximines and residues Asp-261, Asp-294, Gly-363 and Gly-364 (**Supplementary Fig. 7**). With these computational models in hand, free energy perturbation (FEP) calculations^56^ were performed to obtain a quantitative estimate of the relative affinity of the methylsulfoximines **1a** and **1b** for human ASNS. Using a standard thermodynamic analysis (**Fig. 5b**), these calculations provide an estimate of −2.44 kcal/mol for ΔΔG_bind_, meaning that diastereoisomer **1b** has at least 60-fold greater affinity for the enzyme than **1a** at 25 ^o^C (**see Supplementary Information**). This difference is consistent with the expected value based on qualitative arguments that we have presented elsewhere^22^.

### Expression and kinetic characterization of human ASNS variants

We next sought to validate these computational models by examining the effect of site-specific mutations on the ability of ASNS inhibitor **1** to bind to ASNS. Two sets of site-specific mutations were selected on the basis of the intermolecular interactions observed in the computational models of the **1a**/MgPP_i_/ASNS and **1b**/MgPP_i_/ASNS complexes (**Fig. 5c**). Thus, modifying the Glu-364 side chain was anticipated to weaken the binding of diastereoisomer **1a** to the enzyme with little, or no, effect on diastereoisomer **1b**. Similarly, altering the Asp-367 side chain was anticipated to reduce the affinity of **1b** rather than **1a** for the ASNS variant. Using our optimized baculovirus-based protocol, we expressed and purified ASNS variants in which Glu-364 was replaced by alanine (E364A) or glutamine (E364Q), and Asp-367 was replaced by alanine (D367A) or asparagine (D367N). Standard kinetic assays^21,22^ showed that the E364A, E364Q and D367A ASNS variants do not produce pyrophosphate when incubated with L-aspartate and ATP in the presence of ammonium chloride at pH 8.0. The D367N ASNS variant does, however, exhibit ammonia-dependent activity that is substantially reduced relative to that of WT enzyme (**Supplementary Fig. 8**). In agreement with our computational modeling, the time needed for the ASNS inhibitor **1** (at 1 μM concentration) to inhibit ammonia-dependent activity of the D367N ASNS variant is considerably slower than that seen for WT ASNS (**Supplementary Fig. 8**). Interestingly, the presence of **1** also seems to stimulate pyrophosphate production at short times, perhaps because the absence of the Asp-367 side chain leads to an alternate binding mode for **1**. Efforts to characterize inhibitor binding to the three inactive ASNS variants using isothermal calorimetry, however, have proven unsuccessful to date, presumably because complex formation is not completely reversible given slow-onset kinetics and the slow off-rate of the inhibitor in the E*I complex.^21^

### The molecular basis of ASNS inhibitor specificity

With evidence to support the validity of the **1b**/MgPP_i_/ASNS model complex, we next sought to elucidate the molecular basis of the binding selectivity observed for ASNS inhibitor **1**. Superimposing the conserved PP-loop motifs in the three enzymes allowed us to overlay the **1b**/MgPP_i_/ASNS model with the X-ray crystal structures of human argininosuccinate synthetase and GMP synthetase (**Fig. 6a**). Although side chain repositioning was ignored, the superimposed structures provide a qualitative picture of active site similarities and differences that might underlie the binding selectivity of ASNS inhibitor **1** in HCT-116 cell lysates. All three enzymes share a common loop motif for binding MgPPi released during formation of the adenylated intermediate (**Fig. 6a**) and make very similar intermolecular interactions with the AMP moiety of the ASNS inhibitor **1b** (**Fig. 6b**). Differences in inhibitor binding affinity seem to be associated with a “cluster” of negatively charged side chains (Glu-364, Asp-367 and Glu-368) that bind the protonated amino group present in ASNS inhibitor **1** (**Fig. 6c**). This cluster is absent in the active sites of argininosuccinate synthetase (**Fig. 6d**) and GMP synthetase (**Fig. 6e**). This result suggests that second generation ASNS inhibitors must maintain this important set of electrostatic interactions if binding specificity is to be realized. Moreover, given that the conserved residue Glu-364 is required for enzyme activity, it seems unlikely that resistance mutations can occur at this position in the ASNS synthetase active site.

**Fig. 6:**
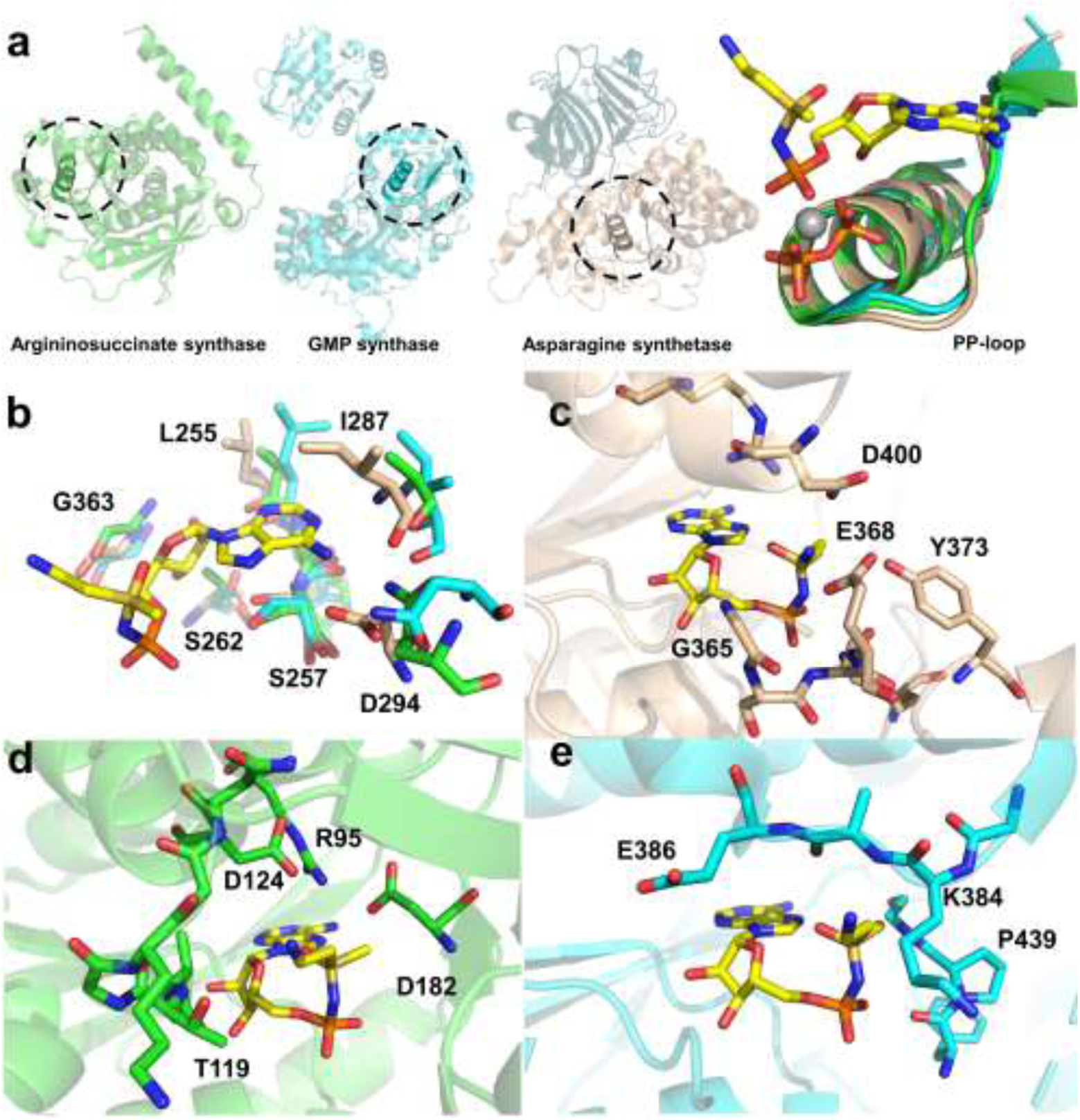
Structural basis for the binding selectivity of ASNS inhibitor 1b. (**a**) Alignment of the conserved SGGxD loops (PP-motifs) in argininosuccinate synthetase (2NZ2)^51^ (green), GMP synthetase (2VXO)^27^ (cyan) and the **1b**/MgPP_i_/ASNS computational model (tan). Circles show the location of the PP-motif in the AMP-forming domain of the three enzymes. Carbon atoms in methylsulfoximine **1b** are colored yellow. Color scheme: N, blue; O, red; P, orange; Mg, grey. (**b**) Superimposition of the homologous AMP-binding sites in argininosuccinate synthetase (C: green), GMP synthetase (C: cyan) and the **1b**/MgPP_i_/ASNS computational model (C: tan) showing the similarity of residues in this region. ASNS residues are labeled using standard one-letter codes, and are numbered from the N-terminal residue (Cys-1). (**c**) Close-up of putative intermolecular interactions between the protonated amino group of **1b** and human ASNS synthetase active site residues (C: tan). ASNS residues are labeled using standard one-letter codes, and are numbered from the N-terminal residue (Cys-1). (**d**) Close-up of argininosuccinate synthetase residues (C:green) surrounding the protonated amino group of **1b** assuming that the ASNS inhibitor binds to the enzyme in a similar pose to that modeled for human ASNS. Argininosuccinate synthetase residues are labeled using standard one-letter codes, and are numbered from the X-ray crystal structure.^51^ (**e**) Close-up of GMP synthetase residues (C: cyan) surrounding the protonated amino group of **1b** assuming that the ASNS inhibitor binds to the enzyme in a similar pose to that modeled for human ASNS. GMP synthetase residues are labeled using standard one-letter codes, and are numbered from the X-ray crystal structure.^27^ The protein orientations in (**c**)-(**e**) are aligned to aid structural comparisons.

## Discussion

Taken overall, our findings establish the feasibility of obtaining ASNS inhibitors that exhibit considerable selectivity when present at low, or sub-, micromolar concentrations in cells despite the existence of other ATP-utilizing enzymes possessing homologous catalytic domains to ASNS. We therefore look forward to the discovery of new small molecule ASNS inhibitors, which can be used (i) to probe the role of L-asparagine production in metastatic progression, and (ii) as agents to control either metastasis and/or tumor growth in animal-based experiments.

## Methods

Methods, including statements of data availability and any associated accession codes and references, are available in the online version of the paper.

## Acknowledgments

We are grateful to Dr. H. Ikeuchi and Professor J. Hiratake (Kyoto) for providing samples of the human ASNS inhibitor **1** used in the functional proteomics experiments. We thank the Diamond Light Source for beam time allocation and beam line staff for assistance with data collection. Mass spectrometric characterizations of human ASNS and the DON-modified variant was performed at the University of Sheffield Biological Mass Spectrometry Facility. This work was supported by Cardiff University (Y.J., P.B. and N.G.J.R.), the National Institutes of Health [R01 GM111695 to Y.T.], the National Science Foundation [MCB-1157688 to Y.T.], and the UK Biotechnology and Biological Sciences Research Council (BBSRC) (BB/I003703/1 to D.W.R.). Financial support for Alexandria Berry was provided by the University of Florida Undergraduate Research Scholar’s Program.

## Author contributions

W.Z., C.B. and S.E.S.: Performed purification and crystallization of human ASNS. W.Z. and C.B.: Performed crystallographic studies and obtained the refined X-ray crystal structure of human ASNS. W.Z.: Performed Dali structural similarity search and analysis. T.I., S.W. and Y.T.: Designed the baculovirus constructs and expressed the recombinant forms of human ASNS. A.R.: Performed the computational modeling studies. B.E.R. and T.K.N.: Devised and performed the functional proteomics experiments. Y.J., P.B. and A.H.B.: Performed kinetic/inhibition assays. W.Z., Y.J., J.W.K., D.W.R., Y.T. and N.G.J.R.: Designed the study. W.Z. and N.G.J.R.: Wrote the manuscript with contributions from all other authors.

## Competing interests

The authors declare no competing financial interests.

## Additional information

Supplementary information is available for this paper. Correspondence and requests for modeling data and materials should be addressed to N.G.J.R.

## References

1. Richards, N.G.J. & Schuster, S.M. Mechanistic issues in asparagine synthetase catalysis. Adv. Enzymol. 72, 145–198.

2. Pui, C.-H., Robison, L.L. & Look, A.T. Acute lymphoblastic leukemia. Lancet 371, 1030–1043 (2008).

3. Yang, H., He, X., Zheng, Y., Feng, W., Xia, X., Yu, X. & Lin, Z. Down-regulation of asparagine synthetase induces cell cycle arrest and inhibits cell proliferation of breast cancer. Chem. Biol. Drug Des. 84, 578–584 (2014).

4. Xu, Y., Ly, F., Zhu, X., Wu, Y. & Shen, X. Loss of asparagine synthetase suppresses the growth of human lung cancer cells by arresting cell cycle at G0/G1 phase. Cancer Gene Ther. 23, 287–294 (2016).

5. Sircar, K., Huang, H., Hu, L., Cogdell, D., Dhillon, J., Tzelepi, V., Efstathiou, E., Koumakpayi, I.H., Saad, F., Luo, D., Bismar, T.A., Aparicio, A., Troncoso, P., Navone, N. & Zhang, W. Integrative molecular profiling reveals asparagine synthetase is a target in castration-resistant prostate cancer. Am. J. Pathol. 180, 895–903 (2012).

6. Knott, S.E, Wagenblast, E., Khan, S., Kim, S.Y., Soto, M., Wagner, M., Turgeon, M.-O., Fish, L., Erard, N, Gable, A.L., Maceli, A.R., Dickopf, S., Papchristou, E.K., D’Santosi, C.S., Carey, L.A., Wilkinson, J.E., Harrell, J.C., Perou, C.M., Goodzari, H., Poulogiannis, G. & Hannon, G.J. Asparagine bioavailability governs metathesis in a model of breast cancer. Nature 554, 378–381 (2018).

7. Hettmer, S., Liu, J., Miller, C.M., Sparks, C.A., Guertin, D.A., Bronson, R.T., Langenau, D.M. & Wagers, A.J. Sarcomas induced in discrete subsets of prospectively isolated skeletal muscle cells. Proc. Natl. Acad. Sci. USA 108, 20002–20007 (2011).

8. Son, J., Lyssiotis, C.A., Ying, H., Wang, X., Hua, S., Ligorio, M., Perrera, R.M., Ferrone, C.R., Mullarky, E., Shyh-Chang, N., Kang, Y., Fleming, J.B., Bardeesy, N., Asara, J.M., Haigis, M.C., DePinho, R.A., Cantley, L.C. & Kimmelman, A.C. Glutamine supports pancreatic cancer growth through a KRAS-regulated metabolic pathway. Nature 496, 101–105 (2013).

9. Hettmer, S., Schinzel, A.C., Tschessaova, D., Schneider, M., Parker, C.L., Bronson, R.T., Richards, N.G.J., Hahn, W.C. & Wagers, A. J. Functional genomic screening reveals asparagine dependence as a metabolic vulnerability in sarcoma. eLife 4, e09436 (2015).

10. Richards, N.G.J. & Kilberg, M.S. Asparagine synthetase chemotherapy. Annu. Rev. Biochem. 75, 629–654 (2006).

11. Cooney, D.A., Jones, M.T., Milman, H.A., Young, D.M. & Jayaram, H.N. Regulators of the metabolism of L-asparagine: A search for endogenous inhibitors. Int. J. Biochem. 11, 519–539 (1980).

12. Cooney, D.A., Driscoll, J.S., Milman, H.A., Jayaram, H.N. & Davis, R.D. Inhibitors of L-asparagine synthetase, *in vitro*. Cancer Treat. Rep. 60, 1493–1557 (1976).

13. Boehlein, S.K., Richards, N.G.J., Walworth, E.S. & Schuster, S.M. Arginine-30 and asparagine-74 have functional roles in the glutamine dependent activities of *Escherichia coli* asparagine synthetase B. J. Biol. Chem. 269, 26789–26795 (1994).

14. Tesson, A.R., Soper, T.S., Ciustea, M. & Richards, N.G.J. Re-visiting the steady-state kinetic mechanism of glutamine-dependent asparagine synthetase from *Escherichia coli*. Arch. Biochem. Biophys. 413, 23–31 (2003).

15. Schnizer, H.G., Boehlein, S.K., Stewart, J.D., Richards, N.G.J. & Schuster, S.M. Formation and isolation of a covalent intermediate during the glutaminase reaction of a class II amidotransferase. Biochemistry 38, 3677–3682 (1999).

16. Boehlein, S.K., Stewart, J.D., Walworth, E.S., Thirumoorthy, R., Richards, N.G.J. & Schuster, S.M. Kinetic mechanism of *Escherichia coli* asparagine synthetase B. Biochemistry 38, 13230–13238 (1998).

17. Koroniak, L., Ciustea, M., Gutierrez, J.A. & Richards, N.G.J. Synthesis and characterization of an *N*-acylsulfonamide inhibitor of human asparagine synthetase. Org. Lett. 5, 2033–2036 (2003).

18. Lücking, U. Sulfoximines: a neglected opportunity in medicinal chemistry. Angew. Chem. Int. Ed. 52, 9399–9408 (2013).

19. Hiratake, J. Enzyme inhibitors as chemical tools to study enzyme catalysis: rational design, synthesis, and applications. Chem. Rec. 5, 209–228 (2005).

20. Schramm, V.L. Enzymatic transition states, transition-state analogs, dynamics, thermodynamics, and lifetimes. Annu. Rev. Biochem. 80, 703–732 (2011).

21. Ikeuchi, H., Ahn, Y., Otokawa, T., Watanabe, B., Hegazy, L.S., Hiratake, J. & Richards, N.G.J. A human asparagine synthetase inhibitor kills asparaginase-resistant MOLT-4 cells. Bioorg. Med. Chem. 20, 5915–5927 (2012).

22. Gutierrez, J.A., Pan, Y.-X., Koroniak, L., Hiratake, J., Kilberg, M.S. & Richards, N.G.J. An inhibitor of human asparagine synthetase suppresses proliferation of an L-asparaginase resistant leukemia cell line. Chem. Biol. 13, 1339–1347 (2006).

23. Patricelli, M.P., Szardenings, A.K., Liyanage, M., Nomanbhoy, T.K., Wu, M., Weissig, H., Aban, A., Chun, D., Tanner, S. & Kozarich, J.W. Functional interrogation of the kinome using nucleotide acyl phosphates. Biochemistry 46, 350–358 (2007).

24. Rosenblum, J.S., Nomanbhoy, T.K. & Kozarich, J.W. Functional interrogation of kinases and other nucleotide-binding proteins. FEBS Lett. 587, 1870–1877 (2013).

25. Rajput, A., Dominguez San Martin, I., Rose, R., Beko, A., LeVea, C., Sharrat, E., Mazurchuk, R., Hoffman, R.M., Brittain, M.G. and Wang, J. Characterization of HCT116 human colon cancer cells in an orthotopic model. J. Surg. Res. 147, 276–281 (2006).

26. LaRonde-LeBlanc, N., Resto, M. & Gerratana, B. Regulation of active site coupling in glutamine-dependent NAD^+^ synthetase. Nat. Struct. Mol. Biol. 16, 421–429 (2009).

27. Welin, M., Lehtiö, L., Johansson, A., Flodin, S., Nyman, T., Trésaugues, L., Hammarström, M., Gräslund, S. & Nordlund, P. Substrate specificity and oligomerization of human GMP synthetase. J. Mol. Biol. 425, 4323–4333 (2013).

28. Sekine, S., Nureki, O., Shimada, A., Vassylyev, D.G. & Yokoyama, S. Structural basis for anticodon recognition by discriminating glutamyl-tRNA synthetase. Nat. Struct. Mol. Biol. 8, 203–206 (2001).

29. Izard, T. The crystal structures of phosphopantetheine adenylyltransferase with bound substrates reveal the enzyme’s catalytic mechanism. J. Mol. Biol. 315, 487–495 (2002).

30. Roionova, I.A., Zuccola, H.J., Sorci, L., Aleshin, A.E., Kazanov, M.D., Ma, C.T., Sergienko, E., Rubin, E.J., Locher, C.P. & Osterman, A.L. Structure of *M. tuberculosis* nicotinate mononucleotide adenylyltransferase. J. Biol. Chem. 290, 7693–7706 (2015).

31. Koizumi, M., Hiratake, J., Nakatsu, T., Kato, H. & Oda, J. Potent transition state analogue inhibitor of *Escherichia coli* asparagine synthetase A. J. Am. Chem. Soc. 121, 5799–5800 (1999).

32. Hinchman, S.K., Henikoff, S. & Schuster, S.M. A relationship between asparagine synthetase A and aspartyl tRNA synthetase. J. Biol. Chem. 267, 144–149 (1992).

33. Lipinski, C.A., Lombardo, F., Dominy, B.W. & Feeney, P.J. Experimental and computational approaches to estimate solubility and permeability in drug discovery and development settings. Adv. Drug Deliv. Rev. 46, 3–26 (2001).

34. Larsen, T.M., Boehlein, S.K., Schuster, S.M., Richards, N.G.J., Thoden, J.B., Holden, H.M. & Rayment, I. Three-dimensional structure of *Escherichia coli* asparagine synthetase B: A short journey from substrate to product. Biochemistry 38, 16146–16157 (1999).

35. Boehlein, S.K., Nakatsu, T., Hiratake, J., Thirumoorthy, R., Stewart, J.D., Richards, N.G.J. & Schuster, S.M. Characterization of inhibitors acting at the synthetase site of *Escherichia coli* asparagine synthetase B. Biochemistry 40, 11168–11175 (2001).

36. Ciustea, M., Gutierrez, J.A., Abbatiello, S.E., Eyler, J.R. & Richards, N.G.J. Efficient expression, purification and characterization of C-terminally tagged, recombinant human asparagine synthetase. Arch. Biochem. Biophys. 440, 18–27 (2005).

37. Imasaki, T., Wenzel, S., Yamada, K., Bryant, M.L. & Takagi, Y. Titer estimation for quality control (TEQC) method: A practical approach for optimal production of protein complexes using the baculovirus expression system. PLoS One 13, e0195356 (2018).

38. Waugh, D.S. An overview of enzymatic reagents for the removal of affinity tags. Prot. Expr. Purif. 80, 283–293 (2011).

39. Handschumacher, R.E., Bates, C.J., Chang, P.K., Andrews, A.T. & Fischer, G.A. 5-Diazo-4-oxo-L-norleucine: reactive asparagine analog with biological specificity. Science 161, 62–63 (1968).

40. Zalkin, H. & Smith, J.L. Enzymes utilizing glutamine as an amide donor. Adv. Enzymol. Relat. Areas Mol. Biol. 72, 87–144.

41. Vagin, A. & Teplyakov, A. Molecular replacement with MOLREP. Acta Crystallogr. Sect. D: Biol. Crystallogr. 66, 22–25 (2010).

42. Brannigan, J.A., Dodson, G, Duggleby, H.J., Moody, P.C.E., Smith, J.L., Tomchick, D.R. & Murzin, A.G. A protein catalytic framework with an N-terminal nucleophile is capable of self-activation. Nature 378, 416–419 (1995).

43. Tesmer, J.G., Klem, T.J., Deras, M.L., Davisson, V.J. & Smith, J.L. The crystal structure of GMP synthetase reveals a novel catalytic triad and is a structural paradigm for two enzyme families. Nat. Struct. Biol. 3, 74–86 (1996).

44. Krahn, J.M., Kim, J.H., Burns, M.R., Parry, R.J., Zalkin H. & Smith, J.L. Coupled formation of an amidotransferase interdomain ammonia channel and a phosphoribosyltransferase active site. Biochemistry 36, 11061–11068 (1997).

45. Boehlein, S.K., Rosa-Rodriguez, J.G., Schuster, S.M. & Richards, N.G.J. Catalytic activity of the N-terminal domain of *Escherichia coli* asparagine synthetase B can be reengineered by single-point mutation. J. Am. Chem. Soc. 119, 5785–5791 (1997).

46. Rognes, S.E. Anion regulation of lupin asparagine synthetase: Chloride activation of the glutamine-utilizing reactions. Phytochem. 19, 2287–2293 (1980).

47. Lomelino, C.L., Andrig, J.T., McKenna, R. & Kilberg, M.S. Asparagine synthetase: Function, structure, and role in disease. J. Biol. Chem. 292, 19952–19958 (2017).

48. Ruzzo, E.K., Capo-Chichi, J.M., Ben-Zeev, B., Chitavat, D., Mao, H., Pappas, A.L., Hitomi, Y., Lu, Y.F., Yao, X., Hamdan, F.F., Pelak, K., Reznik-Wolf, H., Bar-Joseph, I., Oz-Levi, D., Lev, D., Lerman-Sagie, T., Leshinsky-Silver, E., Anikster, Y., Ben-Asher, E., Olender, T., Colleaux, L., Décarie, J.C., Blaser, S., Banwell, B., Joshi, R.B., He, X.P., Patry, L., Silver, R.J., Dobrzeniecka, S., Islam, M.S., Hasnat, A., Samuels, M.E., Aryla, D.K., Rodriguiz, R.M., Jiang, YH., Wetsel, W.C., McNamara, J.O., Rouleau, G.A., Silver, D.L., Lancet, D., Pras, E., Mitchell, G.A., Michaud, J.L. & Goldstein, D.B. Deficiency of asparagine synthetase causes congenital microcephaly and a progressive form of encephalopathy. Neuron 80, 429–441 (2013).

49. Holm, L. and Sander, C. Dali: A network tool for protein structure comparison. Trends Biochem. Sci. 20, 478–480 (1995).

50. Mathews, M.A., Schubert, H.L., Whitby, F.G., Alexander, K.J., Schadick, K., Bergonia, H.A., Phillips, J.D. and Hill, C.P. Crystal structure of human uroporphyrin III synthase. EMBO J. 20, 5832–5839 (2001).

51. Karlberg, T., Collins, R., van den Berg, S., Flores, A., Hammarström, M., Hogbom, M., Schiavone, L.H. and Uppenberg, J. Structure of human argininosuccinate synthetase. Acta Crystallogr. Sect. D: Biol. Crystallogr. 64, 279–286 (2008).

52. Huerta, C., Borek, D., Machius, M., Grishin, N.V. and Zhang, H. Structure and mechanism of a eukaryotic FMN adenylyltransferase. J. Mol. Biol. 389, 388–400 (2009).

53. Bork, P. and Koonin, E.V. A P-loop-like motif in a widespread ATP-pyrophosphatase domain: Implications for the evolution of sequence motifs and enzyme activity. Proteins: Struct. Funct. Genet. 20, 347–355 (1994).

54. Friesner, R.A., Banks, J.L., Murphy, R.B., Halgren, T.A., Klicic, J.J., Mainz, D.T., Rpasky, M.P., Knoll, E.H., Shaw, D.E., Shelley, M., Perry, J.K., Francis, P. and Shenkin, P.S. Glide: A new approach for rapid, accurate docking and scoring. 1. Method and assessment of docking accuracy. J. Med. Chem. 47, 1739–1749 (2004).

55. Ikeuchi, H., Meyer, M.E., Ding, Y., Hiratake, J. and Richards, N.G.J. A critical electrostatic interaction mediates inhibitor recognition by human asparagine synthetase. Bioorg. Med. Chem. 17, 6641–6650 (2009).

56. Abel, R., Wang, L., Harder, E.D., Berne, B.J. and Friesner, R.A. Advancing drug discovery through enhanced free energy calculations. Acc. Chem. Res. 50, 1625–1632 (2017).

